# Highlighting fibroblast plasticity in lung fibrosis: the WI-38 cell line as a model for investigating the myofibroblast and lipofibroblast switch

**DOI:** 10.1101/2023.12.22.572972

**Authors:** Esmeralda Vásquez Pacheco, Manuela Marega, Arun Lingampally, Julien Fassy, Marin Truchi, Kerstin Goth, Lisa Trygub, Marek Bartkuhn, Ioannis Alexopoulos, Ying Dong, Kevin Lebrigand, Andreas Gunther, Chengshui Chen, Cho-Ming Chao, Denise Al Alam, Elie El Agha, Bernard Mari, Saverio Bellusci, Stefano Rivetti

## Abstract

**Background:** Myofibroblasts (MYFs) are generally considered the principal culprits in excessive extracellular matrix deposition and scar formation in the pathogenesis of lung fibrosis. Lipofibroblasts (LIFs), on the other hand, are defined by their lipid-storing capacity and are predominantly found in the alveolar regions of the lung. They have been proposed to play a protective role in lung fibrosis. We previously reported that a LIF to MYF reversible differentiation switch occurred during fibrosis formation and resolution. In this study, we tested whether WI-38 cells, a human embryonic lung fibroblast cell line, could be used to study fibroblast differentiation towards the LIF or MYF phenotype and whether this could be relevant for idiopathic pulmonary fibrosis (IPF).

**Methods:** using WI-38 cells, MYF differentiation was triggered using TGF-β1 treatment and LIF differentiation using Metformin treatment. We analyzed the LIF to MYF and MYF to LIF differentiation by pre-treating the WI-38 cells with TGF-β1 or Metformin first, followed by treatment with Metformin and TGF-β1, respectively. We used IF, qPCR and bulk RNA-Seq to analyze the phenotypic and transcriptomic changes in the cells. We correlated our in vitro transcriptome data from WI-38 cells (obtained via bulk RNA sequencing) with the transcriptomic signature of LIFs and MYFs derived from the IPF cell atlas as well as with our own single-cell transcriptomic data from IFP patients-derived lung fibroblasts (LF-IPF) cultured *in vitro*. We also carried out alveolosphere assays to evaluate the ability of the proposed LIF and MYF cells to support the growth of alveolar epithelial type 2 cells.

**Results:** WI-38 and LF-IPF display similar phenotypical and gene expression responses to TGF-β1 and Metformin treatment. Bulk RNA-Seq analysis of WI-38 and LF-IPF treated with TGF-β1, or Metformin indicate similar transcriptomic changes. We also show the partial conservation of the LIF and MYF signature extracted from the Habermann et al. scRNA-seq dataset in WI-38 cells treated with Metformin or TGF-β1, respectively. Alveolosphere assays indicate that LIFs enhance organoid growth, while MYFs inhibit organoid growth. Finally, we provide evidence supporting the LIF to MYF reversible switch using WI-38 cells.

**Conclusions:** WI-38 cells represent a versatile and reliable model to study the intricate dynamics of fibroblast differentiation towards the MYF or LIF phenotype associated with lung fibrosis formation and resolution, providing valuable insights to drive future research.

**Graphical abstract:** *in vitro* approach using WI-38 cells as a versatile and reliable model to study the MYF or LIF phenotype associated with lung fibrosis formation and resolution observed *in vivo*. WI-38 are providing valuable insights to drive future research on lung fibrosis.

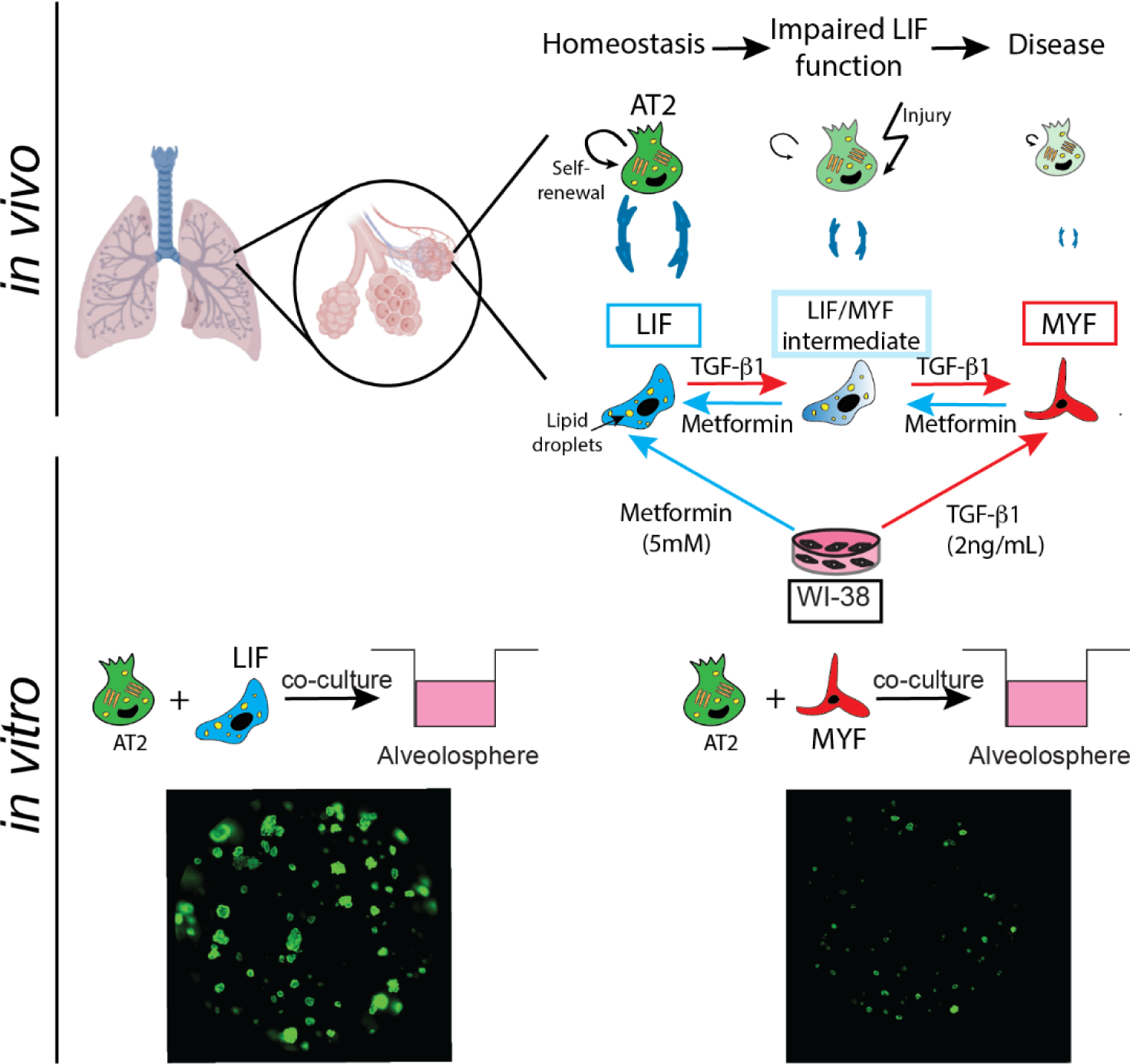

## Introduction

Idiopathic pulmonary fibrosis (IPF) is a debilitating lung disease characterized by excessive tissue scarring, leading to compromised lung function and respiratory failure. Fibrosis disrupts the regular tissue architecture, replacing the normal lung parenchyma with a dense collagen-rich extracellular matrix (ECM) network [1].

Fibroblasts are pivotal players in the intricate web of lung fibrosis pathogenesis. These cells, often described as the architects of the ECM, are indispensable for the structural integrity of tissues. The ECM, a complex network of proteins and carbohydrates, provides mechanical support and signaling cues to cells. When fibroblasts function abnormally or become hyperactive, they can excessively produce ECM components, leading to tissue scarring observed in fibrotic diseases [2]

A particularly intriguing aspect of fibroblasts in the context of lung fibrosis is their plasticity. Under certain conditions, such as tissue injury or specific signaling cues, fibroblasts can undergo differentiation. One well-documented path is their transformation into myofibroblasts (MYFs) [3,4]. Dysregulation of signaling pathways in fibroblasts, driven by factors such as TGF-β1 and others, can trigger pathological fibroblast phenotypes contributing to fibrotic tissue remodeling [5,6].

MYFs are characterized by alpha-smooth muscle actin (ACTA2; α-SMA)[7] expression and robust collagen production. They are often considered the culprits behind the excessive ECM deposition in fibrotic diseases. Their contractile properties, combined with their prolific collagen production, contribute to the stiffening of tissues, a defining feature of fibrotic lesions[8].

On the other hand, fibroblasts can also differentiate into lipofibroblasts (LIFs), a less studied but equally intriguing cell type. LIFs are recognized by their lipid droplet inclusions and have traditionally been associated with roles in lipid storage and metabolism[9]. However, recent research has shed light on their multifaceted functions, especially in lung fibrosis[3]. These cells are not merely passive lipid storages: they actively participate in tissue repair processes and have been implicated in the modulation of fibrosis. Their interactions with other cell types, such as alveolar type 2 cells (AT2), underscore their importance in maintaining lung homeostasis. As our understanding of LIFs deepens, it becomes evident that they play a dual role. While they can be protective and reparative under certain conditions, their dysregulation can also contribute to fibrotic progression [10].

Therefore, fibroblasts, with their potential to differentiate into either MYFs or LIFs, occupy a central position in the narrative of lung fibrosis. Deciphering the signals that dictate their fate and function is crucial for developing therapeutic strategies to halt or even reverse fibrotic diseases[11].

Yet, our mechanistic understanding of the regulation of the MYF or LIF differentiation remains incomplete, mainly due to the limited study models available.

Traditionally, primary lung fibroblast cultures derived from healthy donors and/or IPF patients have been a standard approach to studying fibroblast biology. However, the heterogeneous nature of primary cells, including growth and gene expression variability, presents challenges.

In this study, we propose the use of WI-38 cells, a human embryonic lung fibroblast cell line, as a powerful tool to study fibroblast biology and the transition between lipofibroblasts and myofibroblasts in the context of lung fibrosis. WI-38 cells have been extensively used in vaccine production [12] and studies related to aging including replicative senescence,[13], cancer, and other diseases [14]. They constitute a standard cell line model for cytotoxicity testing [15], due to their genetic stability, ease of handling, and reproducibility of results. However, so far, few studies focused on their differentiation potential [16,17]. Their applicability as a model to study fibroblast differentiation in lung fibrosis has not been fully explored.

First, we compared the results obtained with WI-38 cells with the current primary IPF models. We induced myofibroblast differentiation using TGF-β1 treatment and lipofibroblast differentiation using Metformin treatment. We analyzed the respective phenotypical and transcriptomic changes in WI-38 based on the known markers for myofibroblasts and lipofibroblasts. We also compared their transcriptomic response to the ones elicited by TGF-β1 and Metformin treatment of primary fibroblasts from IPF lungs. We also tested their ability to support alveolar epithelial type 2 cell growth using alveolosphere assays. Moreover, we correlated our *in vitro* transcriptome data from WI-38 cells (obtained via bulk RNA sequencing) with the genetic signature of lipofibroblasts and myofibroblasts derived from the IPF Cell Atlas (http://ipfcellatlas.com/)[18,19] as well as with from our own single-cell transcriptomic data generated on LF-IPF.

The results of our study underscore the potential of WI-38 cells as a versatile and reliable model to study the intricate dynamics of fibroblast differentiation in lung fibrosis, providing valuable insights to drive future research in the field.

## Materials and methods

### Human-derived specimens and cell line

Human primary lung fibroblasts from IPF lungs (referred to as LF-IPF) were obtained from the European IPF registry (eurIPFreg) at the Universities of Giessen and Marburg Lung Center, which is a part of the German Center for Lung Research[20]. Written consent was obtained from each patient, and the ethics committee of Justus-Liebig University Giessen approved the study. The WI-38 cell line (#CCL-75) was purchased from the American Type Culture Collection (ATCC). Cells were amplified, and frozen stocks were made. The cells were used from passage 4 through 7 for the functional assays. Cells were grown at confluence in T75 flasks and split 1:3. Three to four days are required to regain confluency.

### Cell culture

Primary lung fibroblasts derived from 5 IPF patients (LF-IPFs) and the WI-38 cell line were maintained in Dulbecco’s modified Eagle’s medium (DMEM) (Gibco™, cat. nr 21885-025) supplemented with 10% Fetal Bovine Serum (FBS, Gibco™, cat. nr 10270-106) and penicillin/streptomycin (Gibco™, cat. nr 15140-22) at 37°C and 5% CO2. Cells between passages 3 and 7 were used for the experiments. For each experiment, 3 × 10^5^ cells were seeded per well in 6-well plates (Greiner Bio-One). The following day, the cells were starved (0% serum) for 24h and then treated for 96h with 5mM Metformin (Merck, cat. nr M0605000) or 2ng/ml TGF-β1 (Peprotech cat nr 10021). As Controls (Vehicle), the same volume of the TGF-β1 solvent (10mM citric acid, pH 3.0 containing 0.1% bovine serum albumin) was used.

### Transdifferentiation of WI-38 cells and LF-IPFs towards LIF or MYF

Transdifferentiation of fibroblasts to MYF or LIF was determined by immunofluorescence assay and qPCR. The cells (WI-38 and LF-IPFs) were treated with either Vehicle, Metformin or TGF-β1 as described in the previous section. For the immunofluorescence assay aimed at determining MYF transdifferentiation, the following proteins were examined: Actin Alpha 2 (also known as Alpha Smooth Muscle Actin 2, ACTA2), a specific marker for the transition to the myofibroblast, and myosin heavy chain 11 (MYH11), specifically expressed by the activated myofibroblast.

After the treatment, the cells were fixed with 4% formaldehyde for 10min at room temperature. They were then washed with Phosphate Buffered Saline (PBS) and incubated for 1h in a blocking solution (1% Bovine Serum Albumin, 3% goat serum in PBS). For ACTA2 immunostaining, WI-38 or LF-IPF cells were stained with FITC-conjugated mouse monoclonal anti-ACTA2 antibodies at 1:200 dilution (Sigma-Aldrich, cat. nr F3777) overnight at 4°C. For MYH11 immunostaining, both cell types were stained with anti-MYH11 antibodies at 1:100 dilution (Prestige Antibodies power by Atlas Antibodies, Sigma Aldrich, cat. nr HPA015310) overnight at 4°C at room temperature. The following day, the cells were washed with PBS and incubated with secondary antibodies at 1:500 dilution (goat anti-rabbit Alexa 594, Sigma Aldrich,) for 1h at room temperature. After washing, the staining was mounted with DAPI (ProLong™ Gold antifade reagent with DAPI, Invitrogen^TM^ by Thermo Fisher Scientific, cat. nr P36935), and the cells were examined and imaged by fluorescence microscopy (Leica DM5500 B Automated Upright Microscope System). To analyze the presence of lipid droplets, as evidence of differentiation of the cells to a lipofibroblast phenotype, WI-38 and LF-IPFs were treated as described above, and then HCS LipidTOX^™^ green neutral lipid dye at 1:200 dilution (Invitrogen^TM^ by Thermo Fisher Scientific, cat. nr H34475) was added. After incubation at 37°C for 20min, the samples were examined through live imaging and imaged by fluorescent microscopy (Leica DM5500 B Automated Upright Microscope System).

### Staining for lipid-droplet accumulation

To analyze the presence of lipid droplets, as evidence of differentiation of the cells to a lipofibroblast phenotype, LipidTOX green or red neutral lipid dye (Invitrogen^TM^ by Thermo Fisher Scientific, cat. nr H34475) was used (1:200) and cells when alive were placed in the incubation chamber (37 °C and 5% CO_2_) during 20 minutes or 20 min RT when the cells were fixed.

### Immunofluorescence

The cells were fixed in 4% of PFA during 10 minutes, the cells were blocked for 1 hour (3% of BSA, 0.4% Triton X in PBS 1X) at RT, then they were incubated with FITC-conjugated mouse monoclonal anti-ACTA2 (Sigma-Aldrich, cat. Nr. F3777) 1:200 during 1 hour at RT or MYH11 (Prestige Antibodies power by Atlas Antibodies, Sigma Aldrich, cat. nr HPA015310) overnight at 4°C. The following day, for Myh11 immunostaining, the cells were washed with PBS 1X and incubated during 1h at room temperature with Alexa Fluor^TM^ 594 donkey anti-rabbit IgG (H+L) (Sigma-Aldrich, cat. nr A21207) secondary antibody 1:500 was used. After washing, the staining was mounted with DAPI (ProLong™ Gold antifade reagent with DAPI, Invitrogen^TM^ by Thermo Fisher Scientific, cat. nr P36935), and the cells were examined and imaged by fluorescence microscopy (Leica DM5500 B Automated Upright Microscope System).

### RNA extraction and qPCR

The cells were collected in a lysis buffer according to the manufacturer’s protocol, and RNA extraction was performed using the QIAcube® Connect (Qiagen). After quantification using a NanoDrop™ Spectrophotometer 2000/2000c (Thermo Fisher), the RNA was reverse transcribed using the QuantiTect Reverse Transcription kit (Qiagen, cat. nr 205314) and then diluted to a final concentration of 5ng/µL. For qPCR, PowerUp™ Sybr Green Master mix (Applied Biosystems^TM^ by Thermo Fisher Scientific, cat. nr A25742) was used. The primer sequences are listed in Table 1.

**Table 1:**
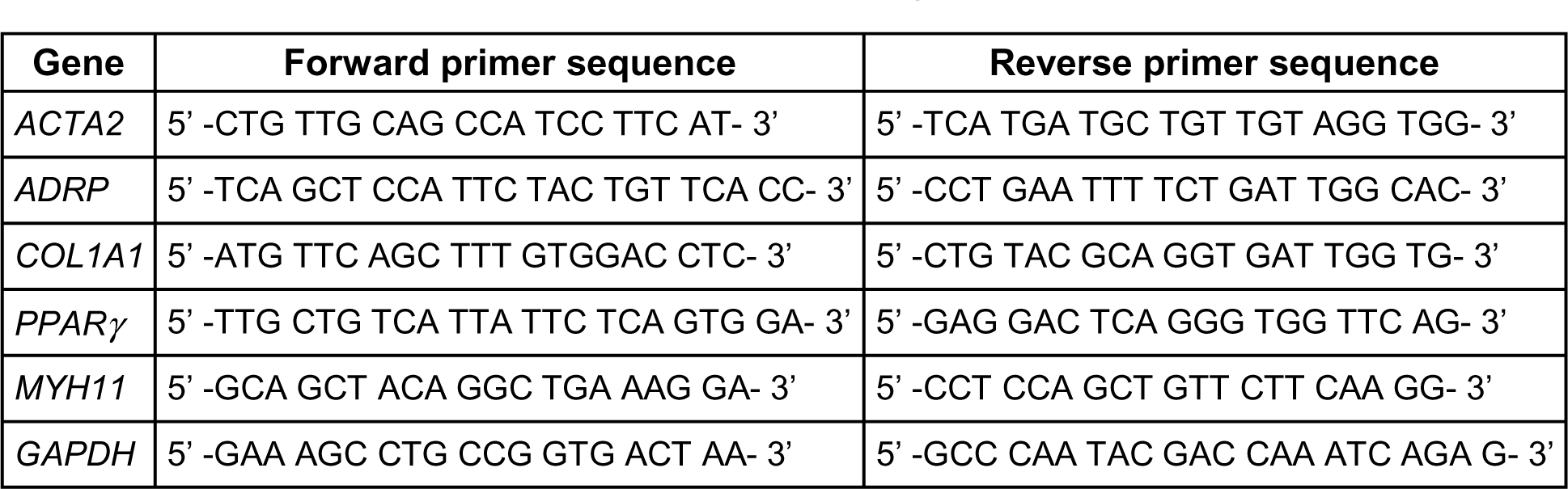
Primer sequences used for qPCR analysis. **Table 1**: List of the forward and reverse primers used in the qPCR analysis: *ACTA2*, *COL1A*, and *MYH11* as MYF markers; *ADPR*, and *PPARγ* as LIF markers. *GAPDH* serves as a housekeeping gene.

**Table 2:** Nucleotide sequence of the 4 HTOs used for Cell Hashing.

|  |
| --- |
| HTO_BC1: AGGACCATCCAA |
| HTO_BC2: ACATGTTACCGT |
| HTO_BC3: AGCTTACTATCC |
| HTO_BC4: TCGATAATGCGA |

### RNA sequencing and bioinformatic analysis

mRNA isolation was performed as described in the previous section. mRNA quality control was assessed. Total RNA was used for library preparation, following the manufacturer’s instructions. All barcoded libraries were pooled and sequenced with 2 × 75 bp paired-end reads on an Illumina NextSeq 550 platform to obtain a minimum of 10 × 10^6^ reads per sample. The generated data were made publicly available. RNA quality was assessed using the Experion RNA StdSens Analysis Kit (BioRad). RNA-Seq libraries were prepared from total RNA using the TruSeq Stranded Total RNA LT kit (Illumina, San Diego, CA, USA), and further analysis was performed as previously published [21]. The statistical analysis of read counts is provided in Table S2 (Statistical analysis of RNA-Seq experiments). RNA-Seq fastq files were checked for quality issues using FastQC (https://www.bioinformatics.babraham.ac.uk/projects/fastqc/).

Trimming and filtering of the reads were performed using Trimmomatic (trimmomatic SE-threads 8 HEADCROP:10 LEADING:3 TRAILING:3 SLIDINGWINDOW:4:15 MINLEN:36). Gene counts were obtained using FeatureCounts [22]. Coverage profiles were created using the Deeptools bamCoverage function with FPKM normalization [23]. Differential gene expression analysis was conducted using the R/Bioconductor package DESeq2 [24]. DESeq2 internal Principal Component Analysis (PCA) was used to visualize the similarity between different samples. Differentially expressed genes were determined based on a false discovery rate (FDR) p-value cutoff of <0.05 and an absolute log2 fold change >1. Heatmaps were generated using the R/Bioconductor package ggplot2 [25] and Venn diagrams were created using the package VennDiagram (https://CRAN.R-project.org/package=VennDiagram). The data generated in this study are available at the GEO repository with the accession number GSE128798. Gene ontology (GO) analyses were performed using the Bioconductor package clusterProfiler[26].

### Viability and cell cycle analysis

Cell viability was evaluated at 2, 24, 48, 72 and 96h using the AlamarBlue^TM^ Cell Viability reagent (Invitrogen^TM^ by Thermo Fisher Scientific, cat nr. DAL1025), following the manufacturer’s protocol. After 96h of treatment, 3000 cells per well were seeded in 96 well plates. AlamarBlue^TM^ Cell Viability reagent (10%) was added to each well. The viability was evaluated by subtracting the average 600nm absorbance values of the cell culture medium alone (used as the blank reference) from the 570nm absorbance values of experimental wells. The absorbance was measured in the Tecan InfiniteM200 plate reader.

### Scratch test for cell migration

WI-38 cells were seeded in a petri dish (Ø 70mm, Greiner Bio-One) at a density of 1.5L×L10^5^ cells per well for each condition previously described. The next day, after 6h of starvation (DMEM-free serum), the cells were treated as previously described. After 96h of treatment, the bottom of each dish was scratched using a sterile pipette tip. After the scratch, the dishes were gently washed with PBS to remove cell debris. After 24h, the washing step was repeated. The cells were maintained in DMEM-free serum at 37 °C and 5% CO_2_ for 96h for the entire assay duration. Images were acquired at 0, 24, 48 and 96h. The percentage of wound closure for each condition (compared to the initial time point) was measured using Image J software with the wound healing size tool [27].

### FACS preparation

B6N.Cg-Tg(Sftpc,-EGFP)1Dobb/J mice (Jackson Lab, strain 028356) were euthanized. The lungs were perfused with 10mL of PBS through the heart’s right ventricle. For the single-cell preparation, lungs were inflated intratracheally with dispase (5U/mL; Corning, cat. nr 354235) and further digested by incubation with 3mL dispase solution (5U/mL) for 20min at room temperature. Lungs were finely chopped and digested in collagenase type IV (Gibco, cat. nr 17104-019) for 30min at 37°C. The cell suspension was passed through 70 and 40µm cell strainers (Greiner bio-one, Easystrainer cat. nr 542070 and 542040, respectively). After washing, the cell suspension was stained with SYTOX^TM^ Red (Invitrogen^TM^ by Thermo Fisher Scientific, cat. nr S34859), and sorting of Sftpc-EGFP^+^ cells was carried out using the BD FACSMelody™ cytometer. Data were analyzed using FlowJo software.

### Alveolosphere assay

21000 sorted Sftpc-EGFP^+^ cells from the adult lungs of B6N.Cg-Tg(Sftpc,-EGFP)1Dobb/J mice and 20000 fibroblast, lipofibroblast or myofibroblasts were resuspended in 50µl of organoid medium (DMEM + 10% FBS + 1% Pen/Streptomycin + ITS 1% (Gibco™, cat. nr 41400-045) + Heparin 0.1% (STEMCELL cat. nr 07980)) plus 50µl of Matrigel (Corning, cat. nr 356231). For each insert, 100µl of the mix was added on top of a 0.4µm insert (Greiner Bio-One), placed in a 24-well plate and incubated for 7min at 37°C. After incubation, 500µl of organoid medium was placed in the lower chamber, and the plate was placed for 14 days in 5% CO_2_ at 37°C. The organoid medium was changed every second day. ROCK inhibitor (10μM, Y27632 STEMCELL cat. nr 72304) was included in the organoid medium for the first 2 days of culture. The percentage of colony-forming efficiency (%CFE) was calculated as the ratio between the number of spheres observed over the initial number of AT2 cells multiplied by 100.

### LF-IPF stimulation and preparation for chromium^TM^ single-cell RNA-seq

LF-IPF (donors #5 and #6) were seeded at 300,000 cells / well in DMEM 10% FCS for 24h. The following day, the cells were starved (0% serum) for 24h and then treated or not for 72h with 5mM Metformin. Following trypsinization, cells from each condition were labelled with in-house Hashtag oligonucleotide (HTO)-coupled antibody (anti-CD90, BD Biosciences, Ref #550402) following the procedure of Stoeckius et al.[28] using the LYNX Rapid Streptavidin Antibody Conjugation Kit (Biorad, Ref #LNK163STR). Briefly, for each condition, 1.10^6^ cells were resuspended in PBS 2% BSA, 0.01% Tween and incubated with 10μL Fc Blocking reagent for 10 minutes at 4°C then stained with 0.5μg of cell hashing antibody for 20 minutes at 4°C. After washing with PBS, 2% BSA, 0.01% Tween, samples were counted and and assessed for single cell separation and overall cell viability (>90%). Samples were then adjusted to the same concentration, mixed in PBS supplemented with 0.04% of bovine serum albumin at a final concentration of 100 cells/μl and pooled sample was immediately loaded onto 10X Genomics Chromium device to perform the single cell capture (3000 cells / condition for a total of 12,000 cells).

### Single-cell RNA-seq data processing

Libraries were prepared as recommended, following the Chromium Next GEM Single Cell 3’ Reagent Feature Barcoding kit (10X Genomics). Libraries were then quantified, pooled and sequenced on an Illumina NextSeq 500. Alignment of reads from the single cell RNA-seq library and unique molecular identifiers (UMIs) counting were performed with 10X Genomics Cell Ranger tool (v3.0.2). Reads of HTOs used for Cell Hashing (BC1-4, see nucleotide sequences below) were counted with CITE-seq-Count (v1.4.2). Counts matrices of total UMI and HTOs were integrated on a single object using Seurat R package, from which the data were processed for analysis. HTOs were demultiplexed with HTODemux in order to assign LF-IPF identity and treatment. Only cells identified as “Singlet” after both demultiplexing and passing quality control thresholds of UMI and mitochondrial content were kept. Differential expression analyses between cells from control and metformin treated LF-IPF were carried out with DESeq2 (v1.30.1). Genes considered as differentially expressed were selected according to an adjusted pvalue threshold of 0.05, obtained with the Wald test and the Benjamini-Hochberg method for multiple tests correction implemented in DESeq2.

### Statistical analysis

Statistical analyses and graph assembly were performed using GraphPad Prism 6 (GraphPad Prism Software). Student’s t-test (unpaired, two-tailed) was utilized to compare the means of two groups, while one-way ANOVA (with post hoc analysis) was used to compare the means of three or more groups. ROUT analysis was performed to assess the presence of outsiders. The corresponding figure legends indicate the number of biological samples (n) for each group and the statistical tests used. Differences in means were considered statistically significant if p < 0.05.

## Results

### Comparison of WI-38 with Primary lung fibroblasts from idiopathic pulmonary fibrosis patients (LF-IPF)

At first, we focused on the behavior of the WI-38 cells and LF-IPF from four different patients in response to the vehicle, TGF-β1 and Metformin treatment for 96h, which induce the fibroblast (FIB), myofibroblast (MYF) and lipofibroblast (LIF) phenotypes, respectively (Figure 1A, B). The cells arising from these treatments, called FIB, MYF1 and LIF1, were analyzed by IF and qPCR.

**Figure 1:**
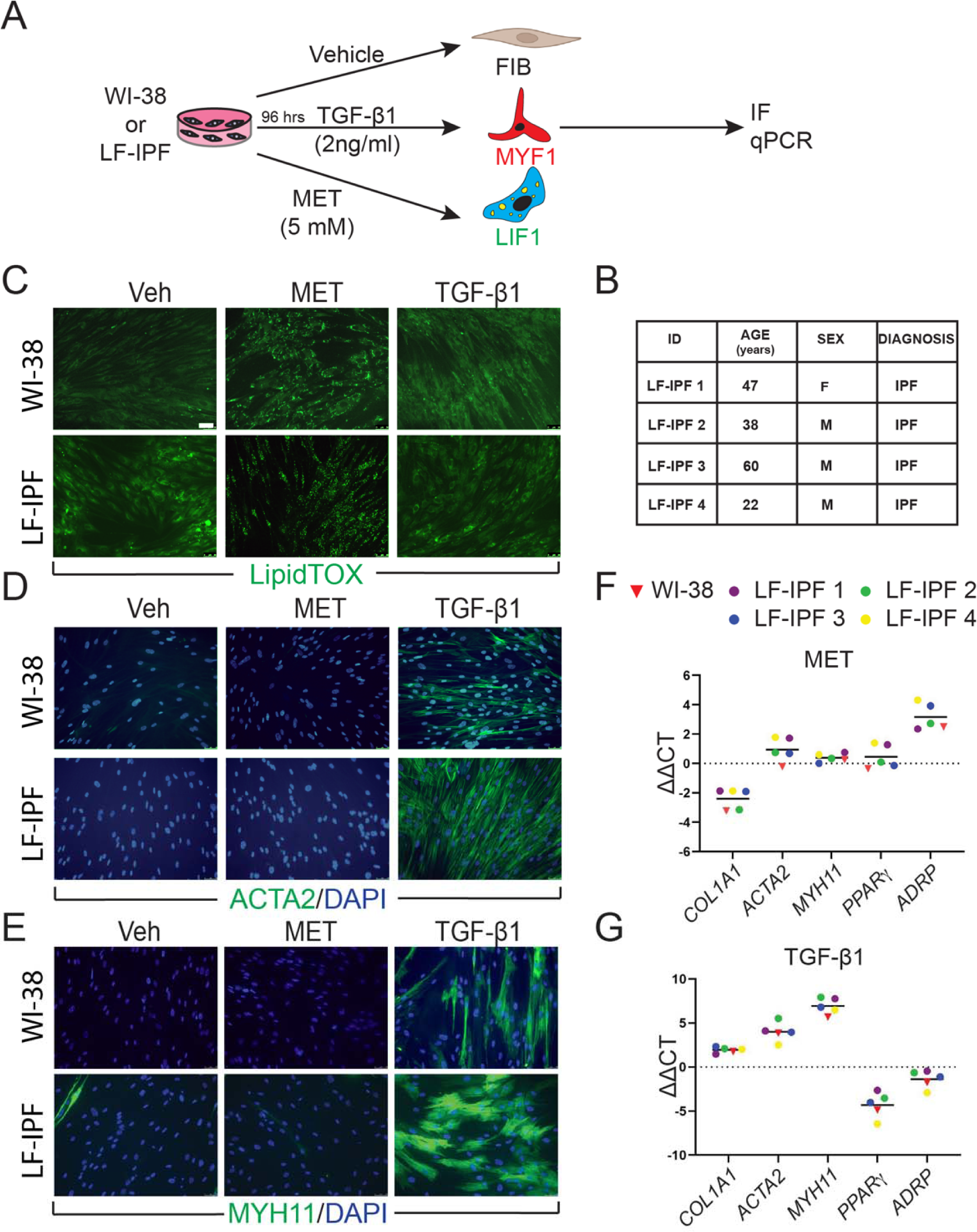
WI-38 and primary lung fibroblasts from idiopathic pulmonary fibrosis patients (LF-IPF) display similar phenotypical and gene expression response to TGF-β1 and Metformin treatment. **A)** Experimental approach. **B)** details on the 4 different LF-IPF used for the study. **C)** IF for LipidTox™. **D)** IF for ACTA2. **E)** IF for MYH11. **F)** qPCR for Metformin vs. vehicle WI-38 and LF-IPF samples. **F)** qPCR for TGF-β1 vs. vehicle WI-38 and LF-IPF samples. Scale bar: 50 μm

The cells were stained with LipidTox™ to detect the presence of lipid droplets, a hallmark of the LIFs. Upon administration, Metformin-treated WI-38 and LF-IPF cells similarly displayed lipid droplet accumulation within the cellular cytoplasm, characteristic of LIF differentiation, while in the cells treated with TGF-β1, the presence of lipid droplets was limited (Figure 1C). The typical markers of MYFs were detected in TGF-β1-treated cells as elevated ACTA2 and MYH11 expression (Figure 1D, E). WI-38 and LF-IPF cells responded similarly to these treatments.

Further examination of gene expression by qPCR also highlighted the remarkable similarity between LF-IPFs and WI-38 (Figure 1F, G). Metformin-treated WI-38 cells showed a significant increase in *Adipose Differentiation-Related Protein* (*ADRP* Aka *PLIN2*) expression (p=0.0024 **), consistent with previous research on LF-IPF (Figure 1F) [29]. Conversely, TGF-β1 treatment triggered an increase in MYF markers (*ACTA2* p=0.0027 **, *COL1A1* p=0.0031 **, *MYH11* p=0.0078 **) in WI-38 cells, mirroring responses seen in primary fibroblasts (Figure 1G)[30]. *PPARγ* expression was also reduced upon TGF-β1 treatment, suggesting an attenuation of the LIF phenotype.

### scRNA-seq analysis of LF-IPF treated or not with Metformin

Next, we evaluated the impact of Metformin treatment on LF-IPF at the single cell level. Two additional independent LF-IPF samples (LF-IPF #5 and LF-IPF #6) were treated with vehicle or 5mM Metformin in SVF-free MEM. After trypsinization, cells from each sample were labeled with a specific HTO barcoded antibody (CD90), pooled, and simultaneously sequenced using droplet based scRNA-seq (10X Genomics Chromium). Overall, we found a balanced representation for each of the four samples, with more than 2000 cells in each condition and a low percentage of doublets and negative singlets (Figure S1). UMAP integrating vehicle- and Metformin-treated LF-IPF for the 2 independent samples considered are shown in Figure 2A, B indicating that virtually all cells from the 2 distinct samples strongly responded to Metformin. Upon Metformin treatment, the expression of *COL1A1* is drastically decreased and the expression of *PLIN2* increased. The Metformin response was highly similar for the two samples indicating that *GDF-15, CSF2* (Growth Differentiation Factor 15, Colony Stimulating Factor 2) and *PLIN2* are among the genes positively regulated (Figure 2C, D), with also a strong conservation of a large subset of significantly upregulated genes when compared to WI-38 stimulated cells (Figure S2). Overall, the global signature of Metformin in LF-IPF was strongly associated with a reduction of extracellular matrix organization and a positive regulation of the inflammatory response, cellular response to lipids and endoplasmic reticulum stress (Figure 2E, F).

**Figure 2:**
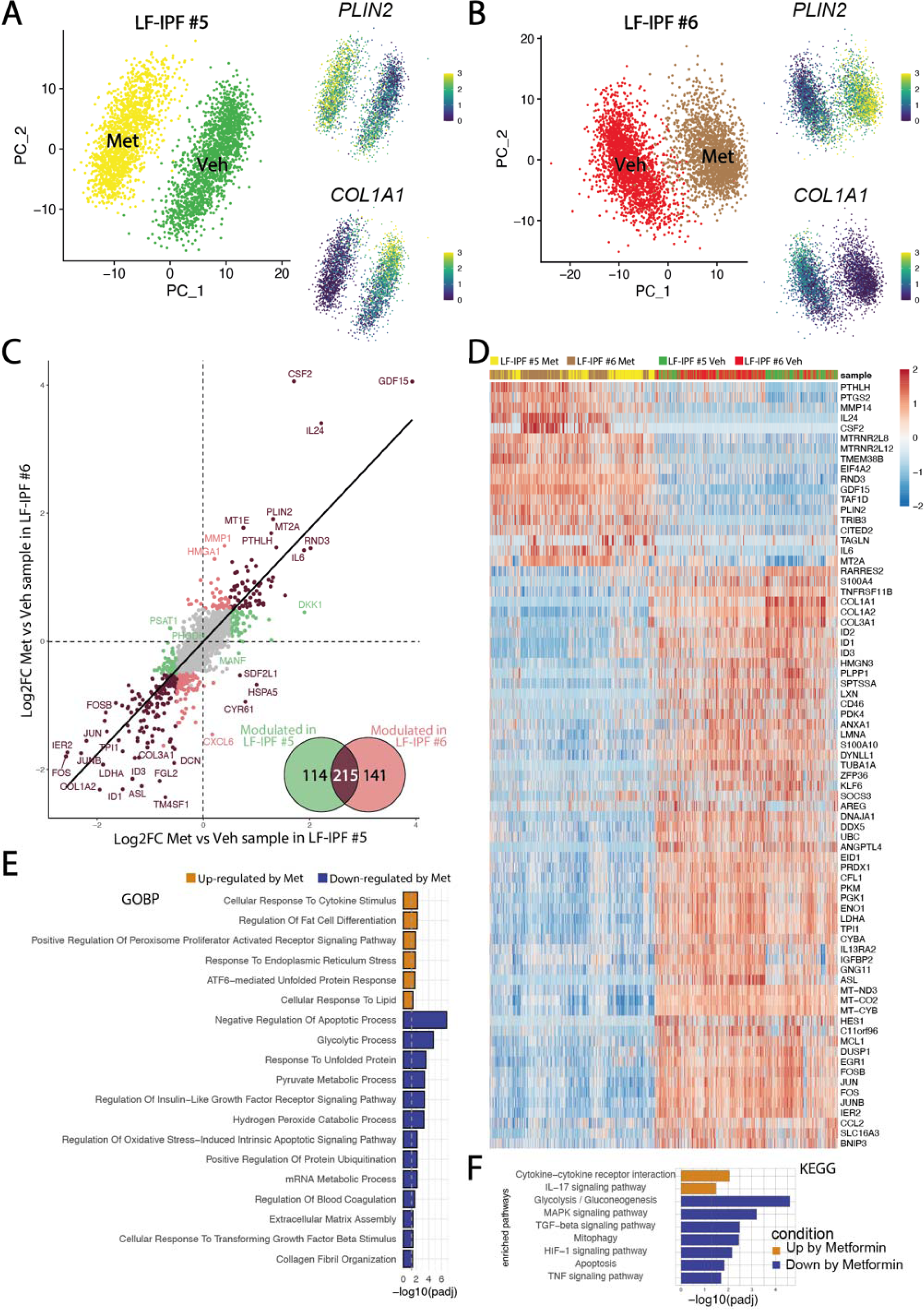
Impact of Metformin treatment on LF-IPF: a scRNA-seq analysis. **A and B)** UMAP integrating vehicle and Metformin treated LF-IPF for the 2 independent samples considered. Please note that the expression of *COL1A1* and *PLIN2* is drastically decreased and increased by Metformin treatment, respectively. **C**) Correlation of the Metformin response in the LF-IPF. The log2 Fold change (Metformin/Vehicle) for each gene is plotted. Black dots correspond to genes significantly modulated in both LF-IPF #5 and LF-IPF #6. Green and red dots correspond to genes significantly modulated in either LF-IPF #5 and LF-IPF #6, respectively. Venn diagram shows the genes modulated by Metformin in the 2 LF-IPF. **D)** Heatmap integrating the 4 samples showing the main gene markers for each condition. **E and F**) Functions and pathway enrichment analysis on differentially regulated genes between Metformin and Control. GOBP: Gene Ontology Biological Processes ; KEGG: Kyoto Encyclopedia of Genes and Genome pathways.

Given the previous observation that the LF-IPF samples were heterogenous in terms of MYF and LIF markers, we carried out a fine clustering of the two LF-IPF samples in control conditions. Figure 3A show the UMAP for the two independent vehicle-treated LF-IPF. Corresponding expression of the *COL1A1* and *PLIN2* on the UMAP suggest the simultaneous presence of MYF and LIF subpopulations in LF-IPF samples. The analysis of the differentially expressed genes in LIF vs MYF subclusters in the two LF-IPF vehicle-treated samples indicated a strong correlation between these subpopulations within the two samples (Figure 3B,C) and revealed, in addition to *PLIN2* the presence of *GDF15* as well as several metallothioneins (*MT1A, MT1E, MT1G, MT1X, MT2A*) in the LIF subpopulation (cluster 2) while genes coding for several collagens (*COL1A1, COL1A2, COL3A1, COL5A2, COL6A3*) were enriched in the MYF population (cluster 0) (Figure 3C). Functional annotation of the 2 clusters confirmed the relevance of these 2 clusters with LIF and MYF, respectively (Figure 3D, E). Finally, Figure S2, shows a 67 gene signature corresponding to the common genes upregulated by Metformin in both WI-38 cells and LF-IPF and found elevated in the LIF subpopulation (cluster 2) identified in basal LF-IPF.

**Figure 3:**
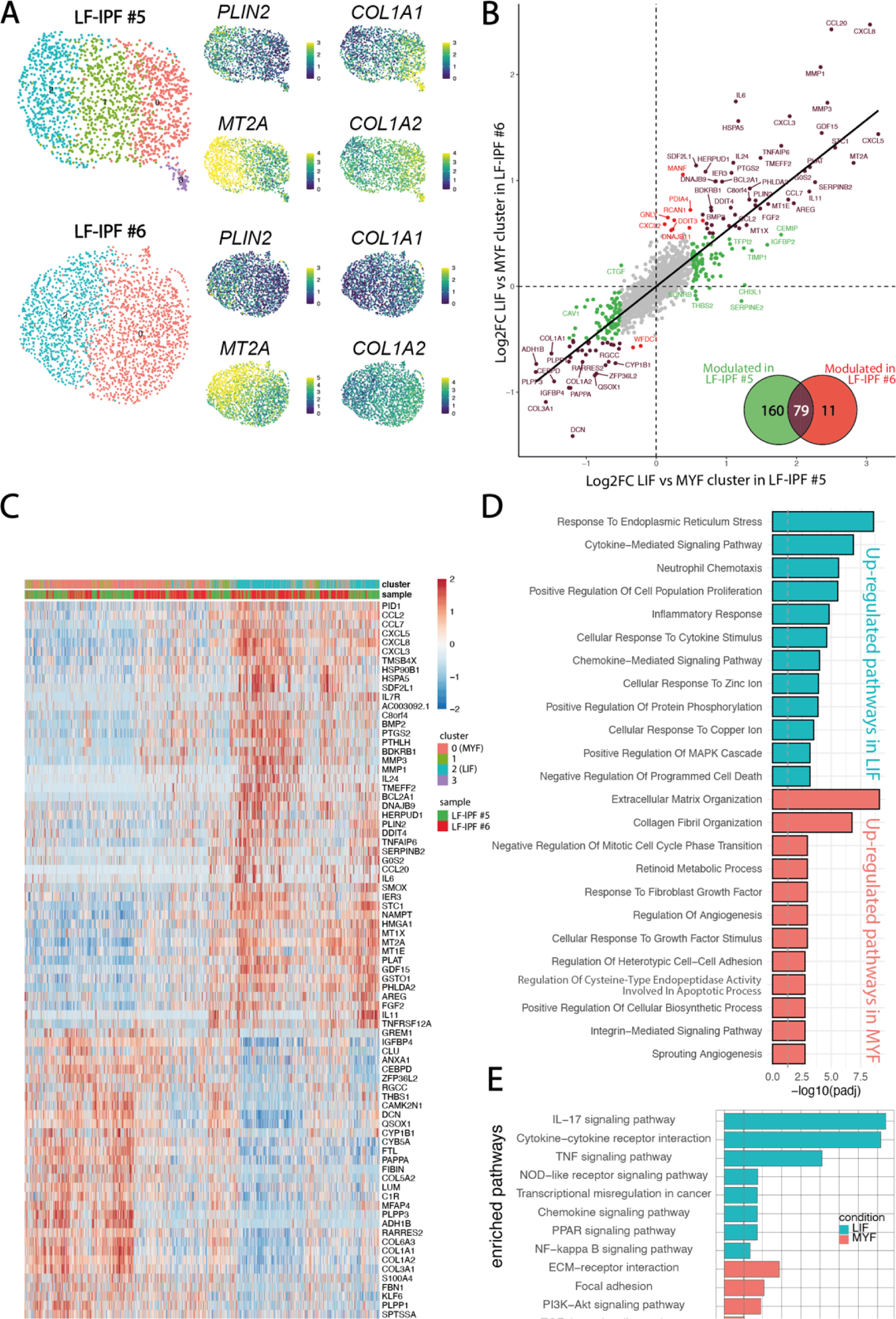
Evidence for presence of LIF and MYF clusters in non-treated LF-IPF samples. **A)** UMAP for the two independent vehicle-treated LF-IPF. Corresponding heatmap and expression of *COL1A1/COL1A2* and *PLIN2/MT2A* suggest the simultaneous presence of MYF and LIF in LF-IPF samples. **B**) Correlation of the LIF vs MYF genes in the 2 LF-IPF. The log2 Fold change (MYF vs LIF) for each gene is plotted. Black dots correspond to genes significantly modulated in both LF-IPF-5 and LF-IPF-6. Green and red dots correspond to genes significantly modulated in either LF-IPF-5 and LF-IPF-6, respectively. The Venn diagram shows the differentially expressed genes between the MYF and the LIF clusters in the 2 LF-IPF. **C)** Heatmap integrating the 2 samples showing the main gene markers for each subclusters. Note the strong similarity between the LIF and MYF clusters for the 2 independent LF-IPF. **D and E**) Functions and pathway enrichment analyses on differentially regulated genes between the LIF and the MYF clusters. GOBP: Gene Ontology Biological Processes ; KEGG: Kyoto Encyclopedia of Genes and Genome pathways.

### Transcriptome Profiling and Comparative Analysis

To test whether the treated- and non-treated WI-38 cells could be separated by their transcriptional changes, principal component analysis (PCA) was applied to the transcriptome profiles of the total genes obtained by bulk RNA-Seq. PCA revealed clear segregation between these three groups, with Metformin-treated cells exhibiting the most dramatic gene expression changes. Additionally, the first two principal components (PC1 and PC2) can separate the three groups, which explains variations among the samples of 60% and 23%, respectively (Figure 4A, B).

**Figure 4:**
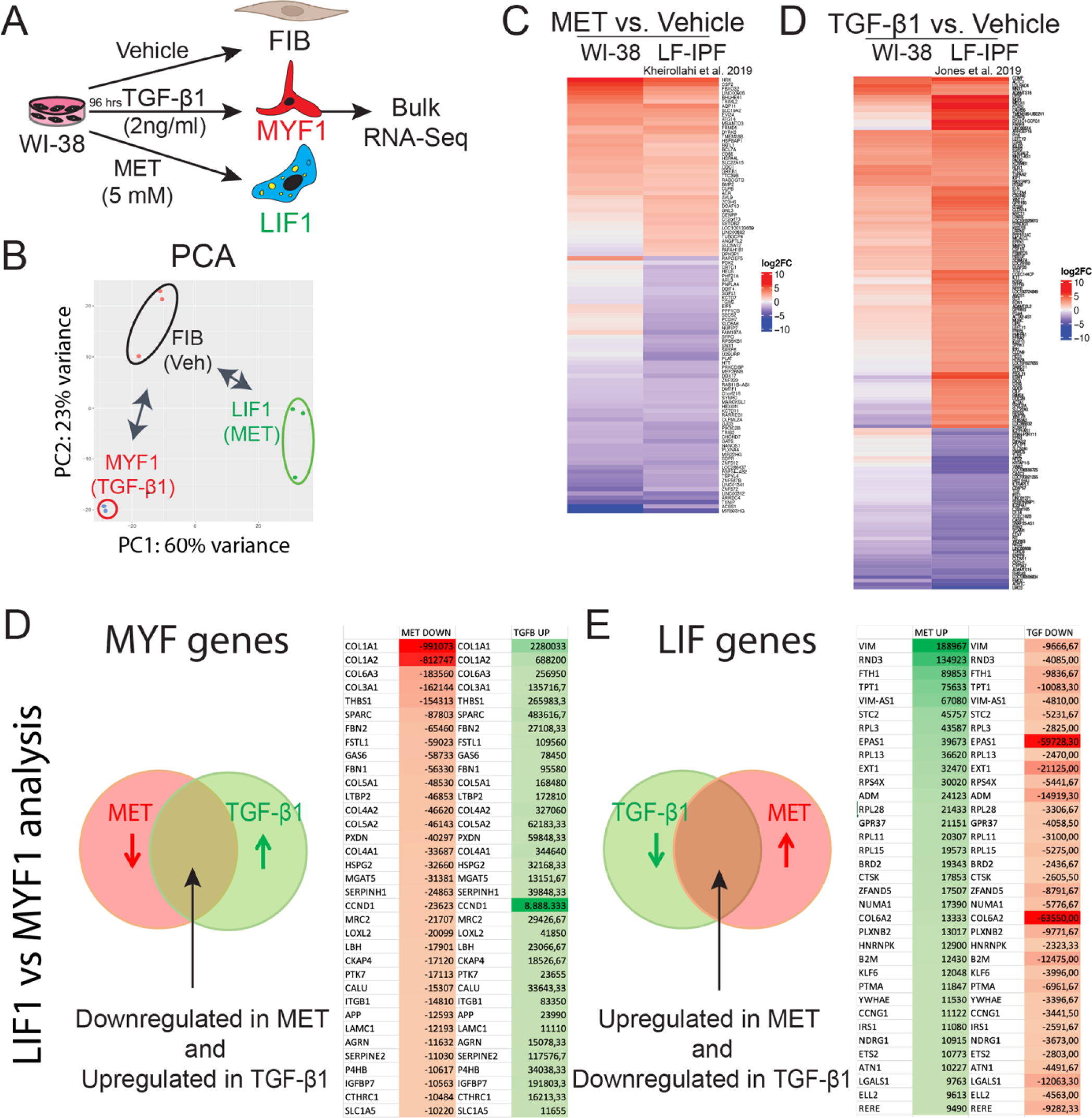
Bulk RNA-Seq analysis of W-I38 and LF-IPF treated with Metformin or TGF-β1 indicate similar transcriptomic changes. **A**) Experimental approach. **B**) PCA analysis. **C**) Comparison of Metformin vs Vehicle treatment for WI-38 and LF-IPF using historical data[29]. **D**) Comparison of TGF-β1 vs Vehicle treatment for WI-38 and LF-IPF using historical data[30]

Subsequent comparison of Metformin-treated WI-38 with previously published data on Metformin-treated LF-IPF showed significant overlap [29]. The derived Heatmap of the top 100 genes revealed the high similarity of the Metformin treated-WI-38 with Metformin treated-LF-IPF (Figure 4C). In agreement with the previous result, the comparison between LF-IPF versus WI-38 treated with TGF-β1 showed a similar expression pattern in the heatmap (Figure 4D).

We also identified the genes simultaneously upregulated by TGF-β1 and downregulated by Metformin (Figure 4D) and selected the top 35 differentially expressed genes based on the absolute gene expression. We propose that these genes are tightly associated with MYF differentiation. Among them, we found well-recognized fibrosis-associated genes such as *COL1A1* and *COL1A2* as well as *Thrombospondin1* (*THBS1*)[31,32] and *Latent Transforming growth factor β binding protein 2* (*LTBP2*)[33], which are encoding for known regulators of latent TGF-β1 activation and critical players in fibrosis. In addition, we identified *Secreted protein acidic and rich in cysteine* (*SPARC*)[34], encoding a glycoprotein involved in fibrosis, as well as *Follistatin-like 1* (*FSTL1*)[35–37], encoding a glycoprotein sequestering inhibitory ligands of TGF-β, and *Serpin H1* (aka *Hsp47*)[38], encoding a collagen specific molecular chaperone associated with increased collagen accumulation. Less known genes involved in fibrosis found in this list are *Peroxidasin* (*PXDN*)[39], *Growth arrest specific gene 6* (*GAS6*)[40,41], *heparan sulfate proteoglycan* (*HSPG2* aka *Perlecan*)[42], *Alpha-1, 6-Mannosylglycopotein 6-Beta-N-Acetylglucosaminyltransferase* (*MGAT5*)[43] as well as *Lysyl oxidase-like 2* (*LOXL2*)[44,45], encoding an enzyme triggering the network of collagen fibers of the ECM, and *Limb-Bud and Heart* (*LBH*)[46], encoding a transcription factor in the WNT/ β-catenin pathway.

Next, we focused on the genes simultaneously downregulated by TGF-β1 and upregulated by Metformin (Figure 4E) and selected the top 35 differentially expressed genes based on the absolute gene expression. We propose that these genes are tightly associated with LIF differentiation. First on this list was *Vimentin* (*VIM*), a gene associated with fibrosis[47]. However, Vim-AS1 was also concomitantly expressed, resulting likely in a low level of Vimentin expression in LIFs. Additionally, we found *Rho family GTPase 3* (*RND3*), encoding a primary antagonist of RhoA activity, which promotes fibrosis[48].

Interestingly, Nintedanib and Pirfenidone upregulate *RND3* expression, suggesting that these 2 drugs act via the inhibition of RhoA activity. We also found *Ferritin Heavy Chain 1* (*FTH1*)[49], which plays a role in iron absorption and transportation and participates in the formation of stored iron. When *FTH1* decreases, the production of stored iron decreases, which leads to the accumulation of intracellular Fe2+, inducing ferroptosis, a cell death mediated by iron-dependent lipid peroxidation. Finally, the presence of *CTSK* (*Cathepsin K*) in this list of LIF genes is consistent with the anti-fibrotic nature of the LIFs, as a high level of *CTSK* is associated with decreased collagen deposition and lung resistance following bleomycin treatment[50].

### Comparative Analysis with Human scRNA-seq Samples

We provided supporting data for using the WI-38 cell line as an alternative model to the LF-IPFs to study LIF and MYF differentiation. To better characterize our model, we extrapolated the LIF and MYF signature obtained from the Habermann scRNA-seq dataset previously published, where 5 Donors and 20 IPF lungs were analyzed (Figure 5A). The expression of the top expressed genes between *PLIN2*+ fibroblasts and Fibroblasts allows to identify 39 genes enriched in *PLIN2+* fibroblasts, including *PLIN2*, *CXCL2*, *MYC* and *GPRC5A* (Figure 5B). The expression of these selected genes representing the LIF signature was then investigated in the Metformin vs. Vehicle bulk RNA-seq data (n=3 independent comparison) from WI-38 cells (Figure 5C). Our results indicated a remarkable upregulation of the LIF signature upon Metformin treatment. The Volcano plot shows the upregulated and downregulated genes in WI-38 cells treated with Metformin using an extended LIF signature containing the top 106 genes expressed (Figure 5D).

**Figure 5:**
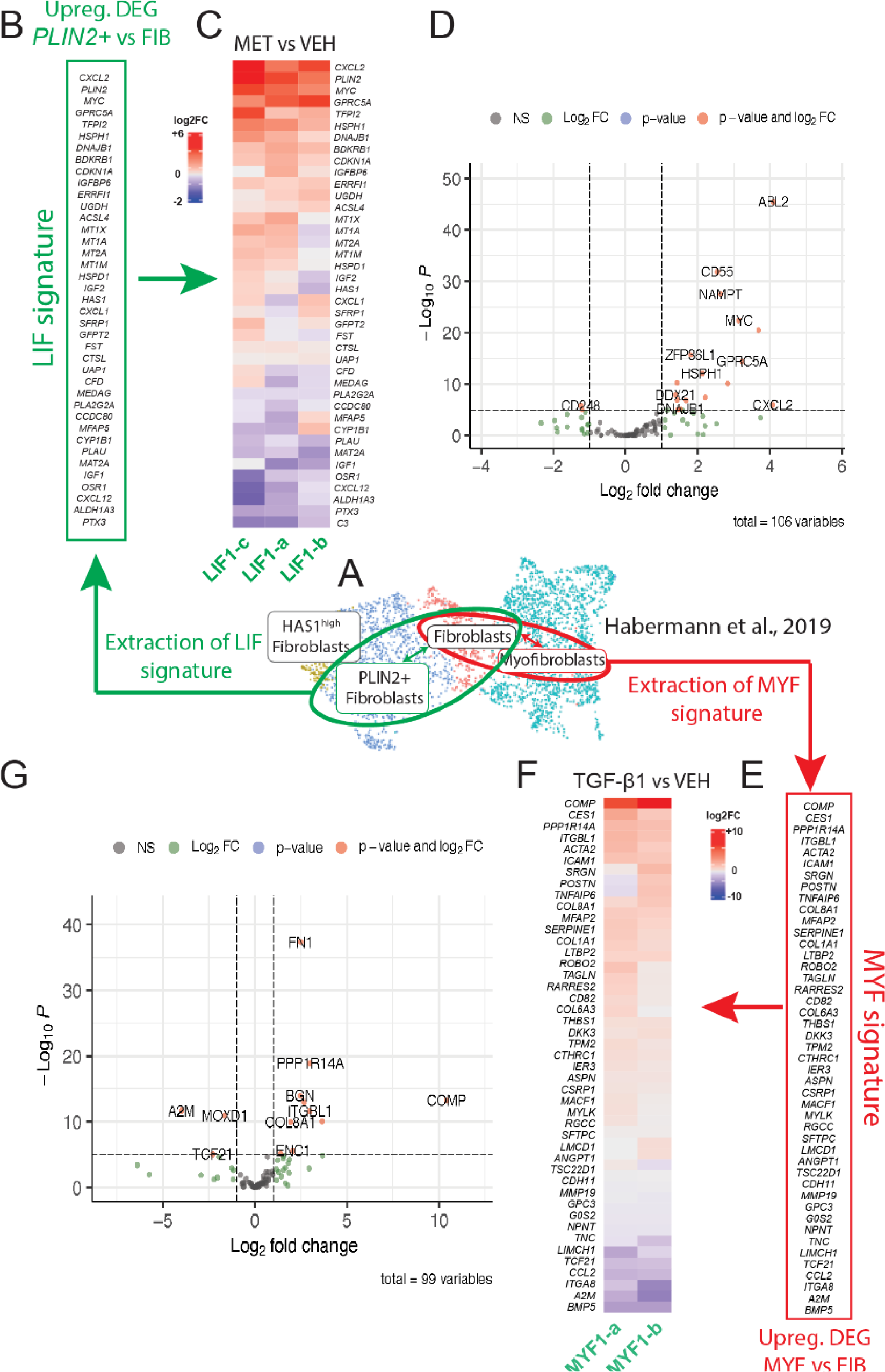
Conservation of the LIF and MYF signature extracted from Habermann et al. scRNA-Seq dataset in WI-38 cells treated with Metformin or TGF-β1, respectively. **A)** UMAP of the Habermann et al., scRNA-seq showing *PLIN2*+ fibroblasts, Fibroblasts and Myofibroblasts. **B)** Extraction of the LIF signature by comparing *PLIN2*+ fibroblasts and Fibroblasts. **C)** Heatmap showing the expression of the LIF signature in Metformin-vs Vehicle-treated WI-38 cells. **D)** Corresponding Volcano plot. **E)** Extraction of the MYF signature by comparing Myofibroblasts and Fibroblasts. **F)** Heatmap showing the expression of the MYF signature in TGF-β1 vs Vehicle-treated WI-38 cells. **G)** Corresponding Volcano plot.

The expression of the top expressed genes between MYFs and FIBs allows us to identify 45 genes enriched in MYFs, including *COMP, CES1, ACTA2, ITGBL1, COL1A1*, and *CTHRC1* (Figure 7E). The expression of these selected genes representing the MYF signature was then investigated in the TGF-β1 vs. Vehicle bulk RNA-Seq data (n=2 independent comparison) from WI-38 cells (Figure 5F). We observed an evident upregulation of the MYF signature in WI-38 cells upon TGF-β1 treatment. The Volcano plot indicates the upregulated and downregulated genes in WI-38 cells treated with TGF-β1 using an extended MYF signature containing the top 99 genes expressed (Figure 5G).

**Figure 7:**
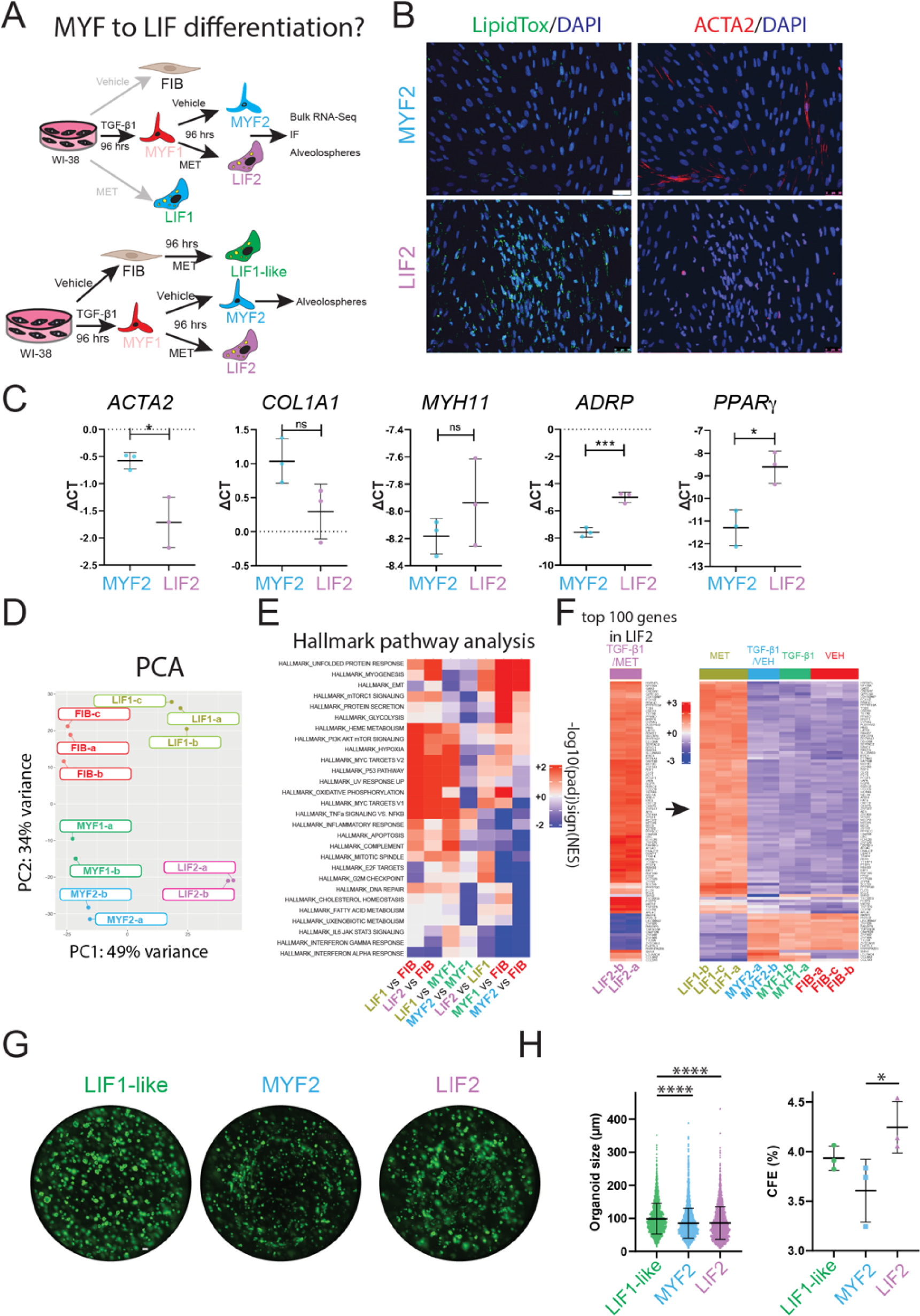
Evidence for MYF to LIF differentiation. **A)** Experimental approach. **B)** IF for LipidTox™ and ACTA2 in MYF2 and LIF2. **C)** qPCR expression of MYF and LIF markers. **D)** PCA graph comparing FIB, LIF1, MYF1, LIF2 and MYF2. **E)** Hallmark pathway analysis. **F)** Heatmap showing the expression of the top 100 regulated genes in LIF2 and the expression of these genes in LIF1, MYF2, MYF1 and FIB. **G)** An alveolosphere assay shows the organoids corresponding to co-culture of LIF1-like, LIF2, and MYF2 with Sftpc^GFP^ cells at day 14. **H)** Quantification of organoid size and colony formation efficiency. Scale bar: 50 μm

For the differential genes in the KEGG pathways enrichment, we used clusterProfiler R package. In the KEGG gene set for LIF differentiation, we observed genes involved in metabolic reprogramming, lipid trafficking, and transcriptional activation. In contrast, for MYF differentiation, we noted an increase in genes related to cytoskeleton reorganization and muscle development (data not shown).

### Functional Assays and Phenotypic Changes

To functionally characterize the capacity of the Metformin- and TGF-β1-treated WI-38 cells to sustain AT2 proliferation, we performed an alveolosphere assay. This assay was carried out by mixing in Matrigel, FIB, LIF1 or MYF1 cells with mouse AT2 cells isolated by FACS using Sftpc^GFP^ lungs (Figure 6A and Figure S3A). Alveolosphere formation is observed within 14 days, as shown in Figure 6B. Quantification of the size of the organoids indicated that LIF1 are significantly supporting the growth of AT2 cells compared to FIB or MYF1 (Figure 6C). No significant difference was observed in the colony forming efficiency between these 3 populations of WI-38 cells (Figure 6D). Taken together, these data are emphasizing the supporting role of LIF on AT2 survival and proliferation, illustrated by the increase in the spheroid size in LIF vs. MYF.

**Figure 6:**
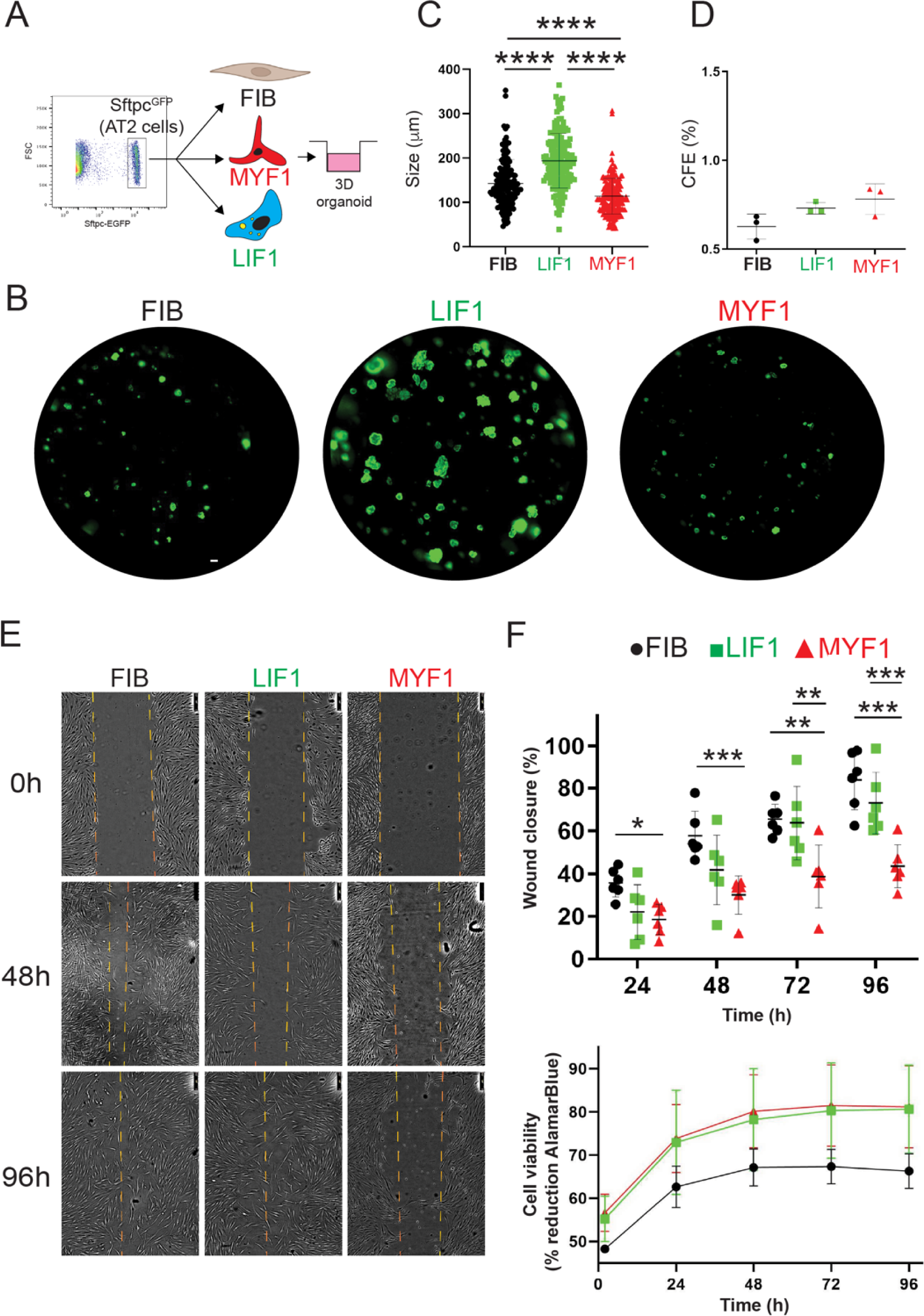
Alveolosphere assays indicate that LIF1 enhance organoid growth while MYF1 elicit organoid growth inhibition. **A)** Experimental approach. **B)** Organoids generated with FIB, LIF1 and MYF1. Quantification of organoid **C)** size and **D)** Colony forming efficiency. **E)** Scratch assay. **F)** Wound closure quantification and cell viability. Scale bar: 250 μm

Additionally, the Metformin-treated cells demonstrated an enhanced migratory capacity in scratch tests, while TGF-β1-treated cells migration capabilities were severely compromised, reflecting a phenotypical feature of fibrotic tissue (Figure 6E, F). These results underscore the practical implications of the LIF and MYF phenotype induced in WI-38. Finally, Alamar Blue viability assay on WI-38 cells treated with Metformin and TGF-β1 revealed no differences between the two groups (Figure 6F), indicating that the treatments did not impact cell viability.

### Evidence for MYF to LIF differentiation

Our results confirm the effectiveness of sequential TGF-β1 and Metformin treatment in inducing the MYF to LIF transition in WI-38 (Figure 7A). After introducing Metformin or vehicle following TGF-β1 treatment, the cells generated after 8 days in culture are called LIF2 and MYF2 (Metformin- and vehicle-treated, respectively). We monitored the expression of MYF and LIF markers in the newly differentiated LIF2 vs MYF2 by IF and qPCR. Our results show a significant decrease in ACTA2 and an increase in lipid droplet accumulation (Figure 7B). Notably, the expression of *ACTA2*, *COL1A1* and *MYH11* was reduced while the expression of *ADRP* and *PPARγ* was increased, correlating with the MYF to LIF transition following Metformin treatment in our cell line.

To better define the nature of these new LIF2 and MYF2, we investigated the global transcriptomic changes occurring in these cells by bulk RNA-Seq. The resulting PCA graph compares LIF2 and MYF2 with the previously generated transcriptomic data for FIB, LIF1 and MYF1 (Figure 7D). Our data indicate that LIF2 are different from LIF1, suggesting that the MYF to LIF differentiation is incomplete. Interestingly, MYF2 appear to display a more significant difference with FIB than MYF1, suggesting a potentially enhanced MYF phenotype.

We also carried out hallmark pathway analysis to compare the different populations to each other (Figure 7E). The comparison of LIF1 vs FIB indicated increased Heme Metabolism, PI3K AKT mTOR signaling, Hypoxia, MYC targets, TNFα signaling vs NFκB as well as decreased Interferon alpha response. Similar regulations for these pathways were observed in LIF2 vs FIB. Direct comparison of LIF2 vs LIF1 indicates decreased TNFα signaling vs NFκB, inflammatory response, apoptosis, complement, IL6/JAK/STAT3, interferon Gamma and Alpha responses. These pathways are also decreased in MYF1 vs FIB and MYF2 vs FIB suggesting that LIF2 represent an intermediate between MYF1 and LIF1. LIF2 are not fully differentiated into a LIF1, still displaying some MYF characteristics.

Additionally, we identified the top 100 genes regulated in LIF2 and examined the expression of these genes in LIF1, MYF2, MYF1 and FIB (Figure 7F). Our results indicated a conservation of these top-regulated genes in LIF1, supporting the MYF to LIF reversion. Moreover, these genes showed opposite regulation in MYF2, MYF1 and FIB, indicating that LIF2 display a different differentiation status compared to the other cell types.

Finally, we assessed the capacity of LIF2, MYF2 and LIF1-like cells to support alveolosphere formation as previously described. (Figure 7G). LIF1-like cells displayed the highest organoid size compared to LIF2 and MYF2. However, the organoid size generated with LIF2 and MYF2 was not different (Figure 7H). Interestingly, LIF2 displayed a significant increase in the CFE compared to MYF2, suggesting that LIF2 confers a survival advantage to the AT2 cells but does not impact their proliferation. Altogether, the difference between LIF2 and MYF2 indicated that LIF2 have partially reversed to a phenotype supportive of alveolosphere formation.

### Evidence for LIF to MYF differentiation

Last, we investigated the LIF to MYF differentiation by pre-treating the WI-38 cells with Metformin followed by TGF-β1 or Vehicle. The cells generated after 8 days in culture are called MYF3 and LIF3. First, we monitored the expression of MYF and LIF markers in LIF3 vs MYF3 by qPCR and IF.

*ACTA2*, *COL1A1,* and *MYH11* expressions were increased, while the expressions of *ADRP* and *PPAR*γ were decreased in MYF3 (Figure 8B).

**Figure 8:**
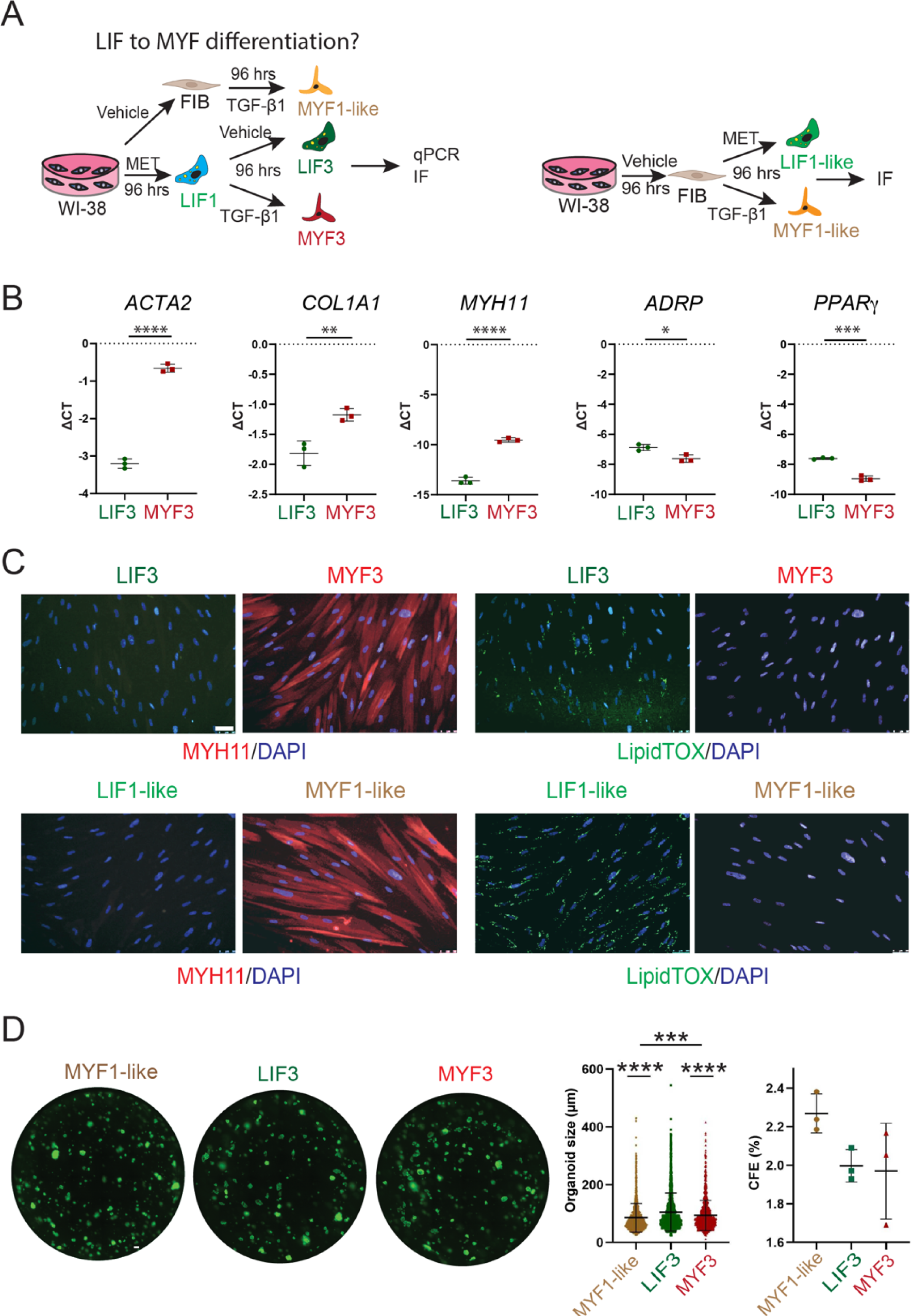
Evidence for LIF to MYF differentiation. **A)** Experimental approach. **B)** IF for LipidTox™ and MYH11 in MYF3 and LIF3. **C)** qPCR expression of MYF and LIF markers. **D)** An alveolosphere assay shows the organoids corresponding to co-culture of LIF1-like, LIF3, and MYF3 with Sftpc^GFP^ cells at day 14. **E)** Quantification of organoid size and colony formation efficiency. Scale bar: 50 μm

A significant decrease in MYH11 and increased in LipidTox™ signal related to the lipid droplets were observed (Figure 8C). LIF1-like and MYF1-like (4 days of starvation followed by either Metformin- or TGF-β1-treatment) were used as controls. Altogether, these results support the LIF to MYF transition in WI-38 cells.

These results strongly validate the WI-38 cell line as a reliable *in vitro* model for studying MYF to LIF differentiation. They exhibit a substantial similarity in their response with the primary human lung fibroblasts.

## Discussion

The heterogeneity of fibroblast subpopulations and their dysregulation in conditions such as idiopathic pulmonary fibrosis (IPF) contribute to tissue structure loss and alveolar reduction. Specifically, myofibroblast activation is crucial in collagen deposition and extracellular matrix remodeling. The MYF population is one of the most extensively studied in the context of fibrosis. However, technical limitations associated with *in vitro* models, such as primary fibroblast cultures, hinder a complete understanding of these cells.

To overcome the limitations associated with primary fibroblast cultures from patients, our study validates the WI-38 cell line, a human embryonic diploid fibroblast cell line, as an *in vitro* model for investigating MYF differentiation upon TGF-β1 treatment. Additionally, we utilized Metformin, an anti-diabetic drug, to induce LIF differentiation in WI-38 cells, in accordance with a previous study [29]. Previous research has shown the ability of these cells to differentiate into LIF and MYF after treatment with parathyroid hormone-related protein (PTHrP) [17,51] and TGF-β1, respectively [16].

WI-38 cells display tremendous technical advantages, specifically in terms of maintenance and culture conditions, genetic manipulation capabilities, and data availability for comparative analysis. One of the most significant advantages of the WI-38 cell line over LF-IPFs pertains to their maintenance and culture conditions. While primary cells often necessitate stringent conditions, such as specialized growth media and carefully controlled environmental factors, WI-38 cells are relatively easy to maintain and culture. The WI-38 cells have a higher propagation capacity, undergoing up to 50 duplications, significantly more than primary fibroblasts. This makes them ideal for long-term experiments, where a continuous supply of cells is essential, thereby minimizing the need for fresh primary cultures and the logistical issues associated with obtaining them.

Another crucial advantage of the WI-38 cell line is its amenability to genetic manipulation, a feature that is often more challenging in primary cultures. The existence of specialized kits designed for the transfection of WI-38 cells further facilitates the process. This not only streamlines the workflow but also opens up new avenues for research, allowing for more complex experimental designs. Such genetic manipulation capabilities are less straightforward in primary cells, which often require more cumbersome techniques and have lower transfection efficiencies. The increasing use and validation of WI-38 cells in multiple studies have led to a wealth of data that can be leveraged for comparative analyses. The consistent genetic background of this cell line makes it easier to interpret and compare data across different studies and laboratories. This is in stark contrast to LF-IPFs, where the genetic variability inherent in primary isolates often leads to batch-to-batch differences, complicating data interpretation and hindering the reproducibility of results. The availability of a standardized, well-characterized cell line like WI-38 enhances the robustness and reliability of data, thus fortifying its utility in large-scale, multi-center studies.

Beyond using WI-38 cells to study MYF differentiation, we report that Metformin-treated WI-38 cells exhibit characteristic features of LIF differentiation, including the accumulation of lipid droplets. LIFs, initially discovered during lung development, were previously regarded as supporting cells for alveolar epithelial type 2 cells (AT2) and a source of lipids for surfactant protein (SFTPC) production. However, recent studies have revealed their active role in supporting AT2 cell proliferation. In the case of lung fibrosis, TGF-β1 induces MYF differentiation in fibroblasts, and LIFs can transdifferentiate into MYFs. However, MYFs cannot support AT2 cell proliferation and differentiation, resulting in alveolarization loss.

In our work, we demonstrated that Metformin treatment triggered the accumulation of lipid droplets in the cytoplasm of both LF-IPFs and WI-38 cells, indicating LIF differentiation. The expression of *adipocyte differentiation-related protein* (*ADRP; PLIN2*) significantly increased upon Metformin treatment, supporting Metformin’s role in lipid accumulation and LIF differentiation.

Concurrently, TGF-β1 treatment amplified MYF-specific markers ACTA2 and MYH11 expression in both LF-IPFs and WI-38 cells, indicating successful induction of the MYF phenotype. Immunofluorescence (IF) and gene expression analysis confirmed the differentiation of WI-38 cells into MYFs and LIFs upon TGF-β1 and Metformin treatment, respectively. Notably, the expression levels of *ACTA2*, *COL1A1*, and *MYH11*, considered markers for MYFs and activated myofibroblasts, were comparable between WI-38 and LF-IPFs. LIF differentiation after Metformin treatment also confirmed the similarity between WI-38 and LF-IPFs, with the accumulation of lipid droplets and the gene expression of *ADRP* and *PPAR*γ overlapping in both models.

Furthermore, sequential treatment of WI-38 cells first with TGF-β1 and then with Metformin led to an exciting reversal of MYF to LIF transdifferentiation, underscoring the pivotal role of Metformin in this cellular reprogramming (for a summary of all the experimental conditions see Figure S3B).

Comparison between scRNA-seq data from human samples and our in vitro model further demonstrated the effectiveness of this approach. The gene expression signatures observed in LIF and MYF cells in our *in vitro* WI-38 models showed a remarkable overlap with human lung samples. This further validates our model and, importantly, the therapeutic potential of Metformin.

Interestingly, our transcriptome analysis revealed a profound impact of Metformin treatment on the gene expression of WI-38 cells. The significant enrichments in the KEGG pathways highlighted Metformin’s effects on metabolic reprogramming and lipid trafficking. However, the exact molecular mechanism by which Metformin achieves this remains a subject for future investigation.

In the organoid formation assay, Metformin-treated WI-38 cells supported alveolosphere formation more effectively, indicating that the LIF phenotype induced by Metformin could significantly impact lung tissue repair. However, fibroblast viability was comparable among all the groups, demonstrating that these therapeutic alterations do not compromise the fundamental properties of the cells.

In conclusion, our results provide compelling evidence of Metformin’s role in lung fibroblast differentiation and its therapeutic potential. The high comparability between the WI-38 cell line and LF-IPFs will allow further study of LIF and MYF differentiation and explore potential therapies for conditions such as pulmonary fibrosis. In addition, our findings emphasize that while Metformin’s role in the lung is still not fully understood, it offers an exciting possibility as a therapeutic tool. The substantial evidence gathered through the WI-38 cell model provides a robust foundation for further exploration of Metformin’s potential roles in lung repair and fibroblast differentiation.

Furthermore, the WI-38 cell line offers compelling advantages over primary fibroblasts, making it a valuable resource for studies focused on lung homeostasis and disease. Its ease of maintenance, higher propagation capacity, and capability for genetic manipulation provide researchers with a versatile and reliable model system. Additionally, the wealth of comparative data available for WI-38 cells enhances its utility, making it a preferred choice for basic and applied research in lung fibroblast differentiation and fibrotic lung diseases. Therefore, our study strongly advocates for the broader adoption of the WI-38 cell line in future research endeavors.

## Acknowledgements

The authors thank the technical support of the UCA GenomiX platform of the University Côte d’Azur. Vasquez-Pacheco E. thanks CONAHCYT for the doctoral fellowship (CVU 768005).

## Contributions

EVP, MM, AL, KG, LT, YD carried out the experiments. AG, CC, CMC, DAA and EAE contributed to the analysis of the results. JF, MT, MB, KL contributed to the bio-informatic analyses. IA contributed to the imaging. BM, SB and SR designed the project, contributed to the analysis of the results, contributed to the bio-informatic analyses and wrote the paper.

## Figures

**Figure S1:**
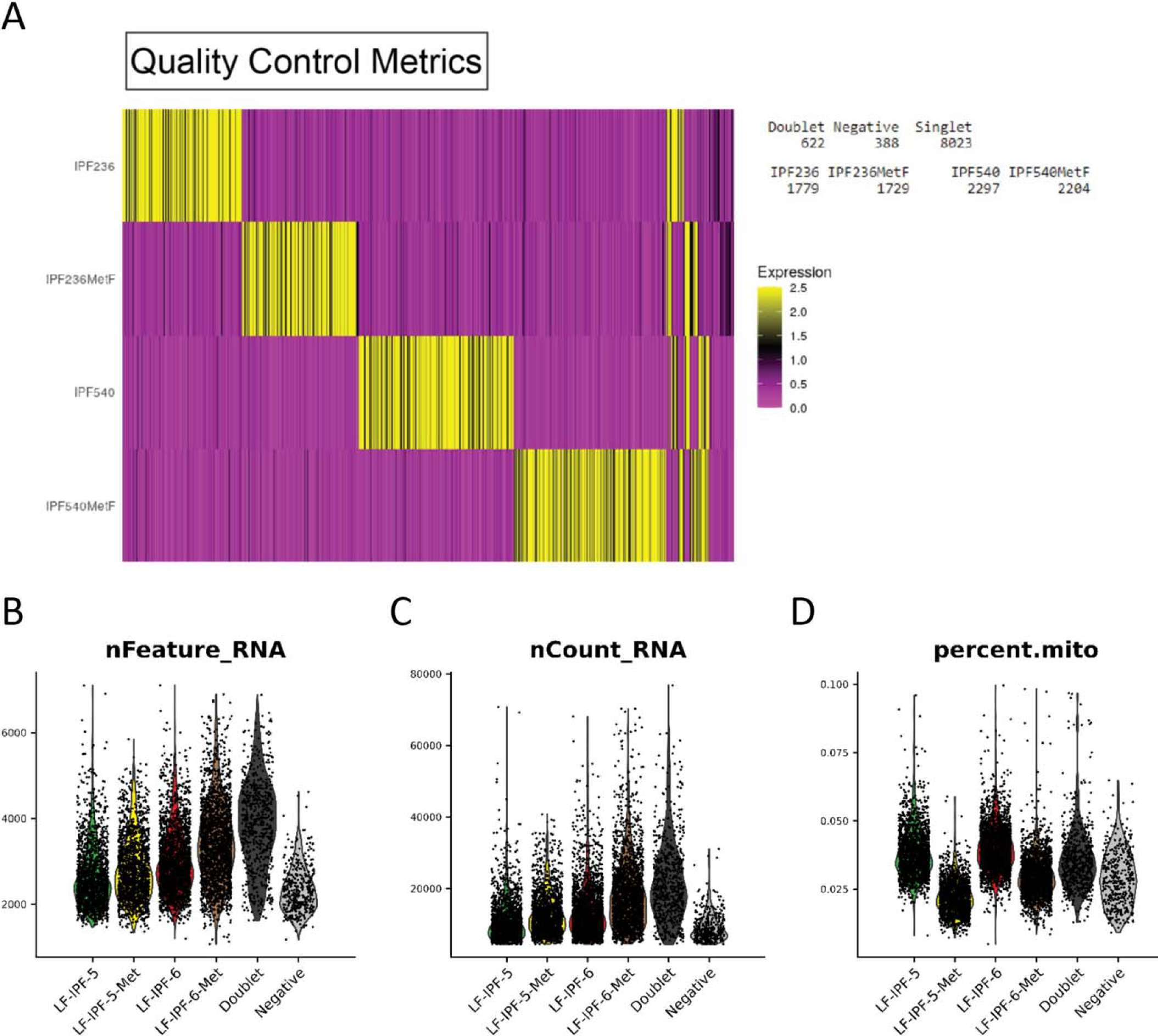
Quality control metrics of the scRNA-seq experiment. **A)** Heatmap of HTO counts in each cell, labelled according to assigned experimental condition after demultiplexing. B-D) Violin plots showing the number of genes detected in each cell (B), the number of UMI per cell (C) and the percentage of cell reads originating from the mitochondrial genes (D).

**Figure S2:**
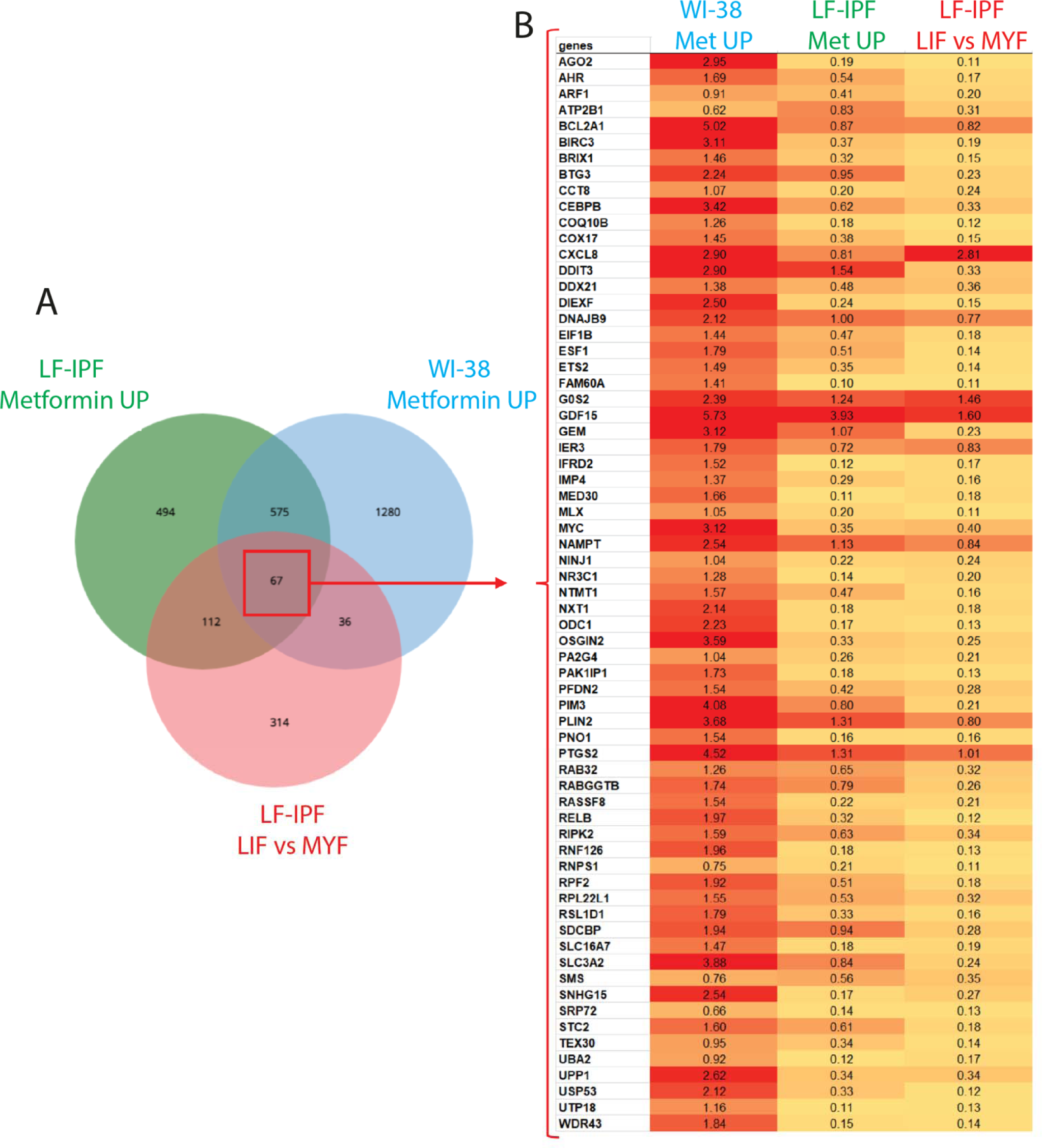
Comparison of the LIF signature in WI-38 treated or not with Metformin, LF-IPF Metformin vs Control (scRNA-seq see Figure 2) and between the proposed LIF vs MYF clusters in non-treated LF-IPF (scRNA-seq Figure 3). **A)** Venn diagram showing the indicated comparison (left) and B) the heatmap of the identified 67 genes common to the 3 comparisons. The log2 ratio for each condition is indicated in the table. The intensity of the red is directly proportional to the expression level.

**Figure S3:**
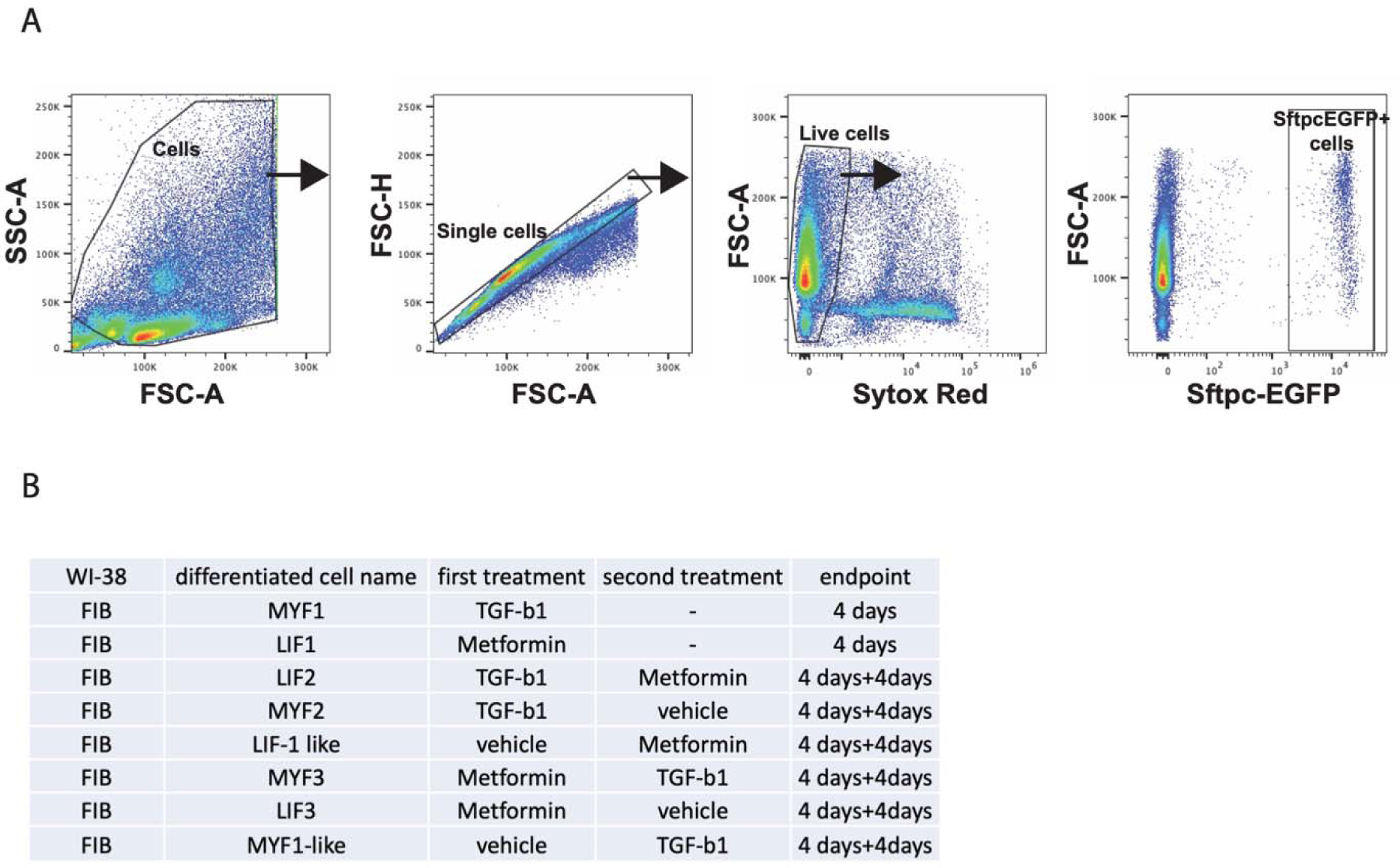
**A)** Gating strategy for the isolation of GFP positive cells from adult Sftpc^GFP^ mouse lungs . **B)** Abbreviation used for the different culture conditions

